# Variance partition reveals contrasting random effect contributions on the density and species composition of malaria-transmitting mosquitoes

**DOI:** 10.1101/2025.07.09.663957

**Authors:** Tin-Yu J. Hui, Patric Stephane Epopa, Abdoul Azize Millogo, Franck A. Yao, Dao Koulmaga, Florian Noulin, Abdoulaye Diabate, Austin Burt

**Author notes:** Corresponding author: Tin-Yu Hui < >.

## Abstract

Spatial-temporal variations exist in the density and species composition of malaria-carrying mosquitoes, which will in turn influence the transmission of the disease. While there has been extensive research on the seasonality and other main drivers of the vector populations, the heterogeneity induced by random effects is just as important but did not quite attract the same attention. To investigate the relative contributions of the between-house, between-village, and between-year variations, as well as other house-level covariates such as inhabitant number and bed net usage, intensive Pyrethroid Spray Catches (PSC) sampling was conducted across a 60-month period between 2012-2019 from four villages in the Sudano-Sahelian region of Burkina Faso.

For mosquito density, measured by counts of female *Anopheles gambiae s.l*., our modelling showed that the between-house variation was the largest component, followed by the between-year then between-village variation, after accounting for seasonality and other covariates. Density increased with the number of inhabitants within a household but was uncorrelated with bed net usage. A subset of female mosquitoes was genotyped for species identification, and the composition of *An. coluzzii* and *An. gambiae*, the two dominant vectors in the region, varied hugely across villages without a clear seasonal trend. The between-village variance contributed up to 76% of the total random variation, followed by the between-year variance. The between-house variation was estimated to be statistically insignificant. Neither household size nor bed net usage had any impact on species composition.

In short, the relative importance of the random components in mosquito density was in the reverse order from species composition. The estimates and relative strengths help parameterise potential field trials for novel vector control programmes and monitoring.

## Introduction

Despite the continued effort and investment in malaria control programmes, regions in sub-Saharan Africa remain disproportionally affected by this vector-borne disease. Harrowing figures of 263 million cases and over half a million deaths are still being reported on the continent, accounting for about 95% of the global malaria burden [1]. Recent modelling predicted that the current intensity and range of interventions (such as long-lasting insecticidal nets and indoor residual spraying) can at best maintain the status quo but are unable to drive malaria towards eradication [2]. In fact, the recent increase in malaria incidence coincides with the vectors’ resistance to insecticides [3, 4], highlighting the need for developing complementary cost-effective and sustainable novel technologies for malaria control. One example is the Sterile Insect Technique (SIT), which has previously been implemented by the release of radiation-sterilised individuals across multiple insect species that are deemed harmful [5]. SIT can also be achieved via Genetic Modification (GM) of the vectors, such as by engineering a transgene into the males of *Aedes aegypti* for dengue control [6]. Beyond SIT, there has been extensive theoretical and laboratory-based research on developing gene drive systems for *Anopheles gambiae s.l*., a species complex whose members contain the primary malaria vectors in sub-Saharan Africa. Gene drive allows the transgene to be passed on to offspring and beyond in a super-Mendelian manner to achieve longer-term population suppression or replacement [7, 8].

For contagious or vector-borne diseases, the basic reproductive number *R*_0_ is the most important metric to quantify their transmission, with *R*_0_ < 1 being a requirement for eradication. In malaria, *R*_0_ is generally defined as the expected number of persons to be infected one generation after the malaria parasite was introduced, or equivalently the number of secondary cases [9]. As the life cycle of the malaria parasite consists of both mosquito and human phases, complex parameterisation arises when characterising the host-parasite-vector interaction [10]. The entomological inoculation rate (EIR), for example, connects the sporozoite rate (i.e., the prevalence of the parasite in the salivary glands of female mosquitoes), and human biting rate (HBR) per night, which has the mosquito density per person embedded [11]. Under the classical models *R*_0_ and vectorial capacity are functions of these entomological rates at equilibrium. To reduce the intensity of transmission, most vector control programmes aim to supress factors concerning the vectors’ density or competence [9].

Malaria transmission is never uniform across space and time. Previous estimates of *R*_0_ varied hugely from ∼1 to >1000 across Africa [9], and mosquito density fluctuates across seasons often by orders of magnitude [12]. In fact, heterogeneity exists on almost all levels: from mosquito density and species distribution across countries, exposure among households within the same village, down to the variable susceptibility of individuals. While earlier works have focused on identifying the main drivers (such as environmental and climatic factors) associated with the trends in malaria vectors and cases [13], it is equally important to quantify the unexplained portion of the variation. Through the extensive mosquito collection survey (see below) this study has the following aims: 1) To estimate the random or unexplained variance components concerning mosquito density and species composition at different spatial-temporal levels, in particular, the between-village, between-house, and between-year variation, after accounting for seasonality and other fixed covariates. 2) To discuss how these random components and their relative strengths impact vector density and malaria transmission. 3) To use these analyses to provide recommendations on the potential design and sample size considerations for clustered randomised controlled trial (cRCT) under various intervention scenarios.

## Sampling

Mosquitoes were sampled in the houses from four villages in Burkina Faso’s Sudano-Sahelian: Bana Market, Bana Village, Soukroudingan, and Pala (Figure 1), where morphologically identical *An. gambiae (s.s.), An. coluzzii*, and *An. arabiensis* are the primary vectors. The first three villages are located to the west of Bobo-Dioulasso, where our research base situates, while Pala is on the opposite direction from the rest [14]. Bana’s two collection sites, Bana Market and Bana Village, are separated by a small semi-permanent river with different human activities [15]. Two phases of collection were conducted: from July 2012 to June 2014 and throughout 2017-2019, covering a total of 60 months. Monthly collection was scheduled aiming to sample 20 houses per village per month. Houses were chosen randomly from a pool of known houses based on local information, from which half of the houses were repeatedly visited over time (fixed houses), while the remaining half were further randomised each month (random houses). For each village, we attempted to sample all houses in the same or consecutive days to minimise the effect of nuisance abiotic factors. For Pala and Souroukoudingan, the sampling interval switched to once every two months in the second phase (Figure 2). Sampling took place after sunrise via indoor pyrethroid spray catches (PSC), using a well-known and locally-market available insecticide spray Kaltox® Paalga (Sphyto company, Burkina Faso). All mosquitoes were morphologically identified in the field [16]. The targeted organisms *An. gambiae s.l*. were preserved in 80% ethanol for further analysis. During the second phase of collection PCR was performed on a subset of the preserved samples for species identification, such that the composition of the species (within *An. gambiae s.l*.) per visit was also known. The genotyping was based on the detection of the SINE 200x locus [17].

**Figure 1.**
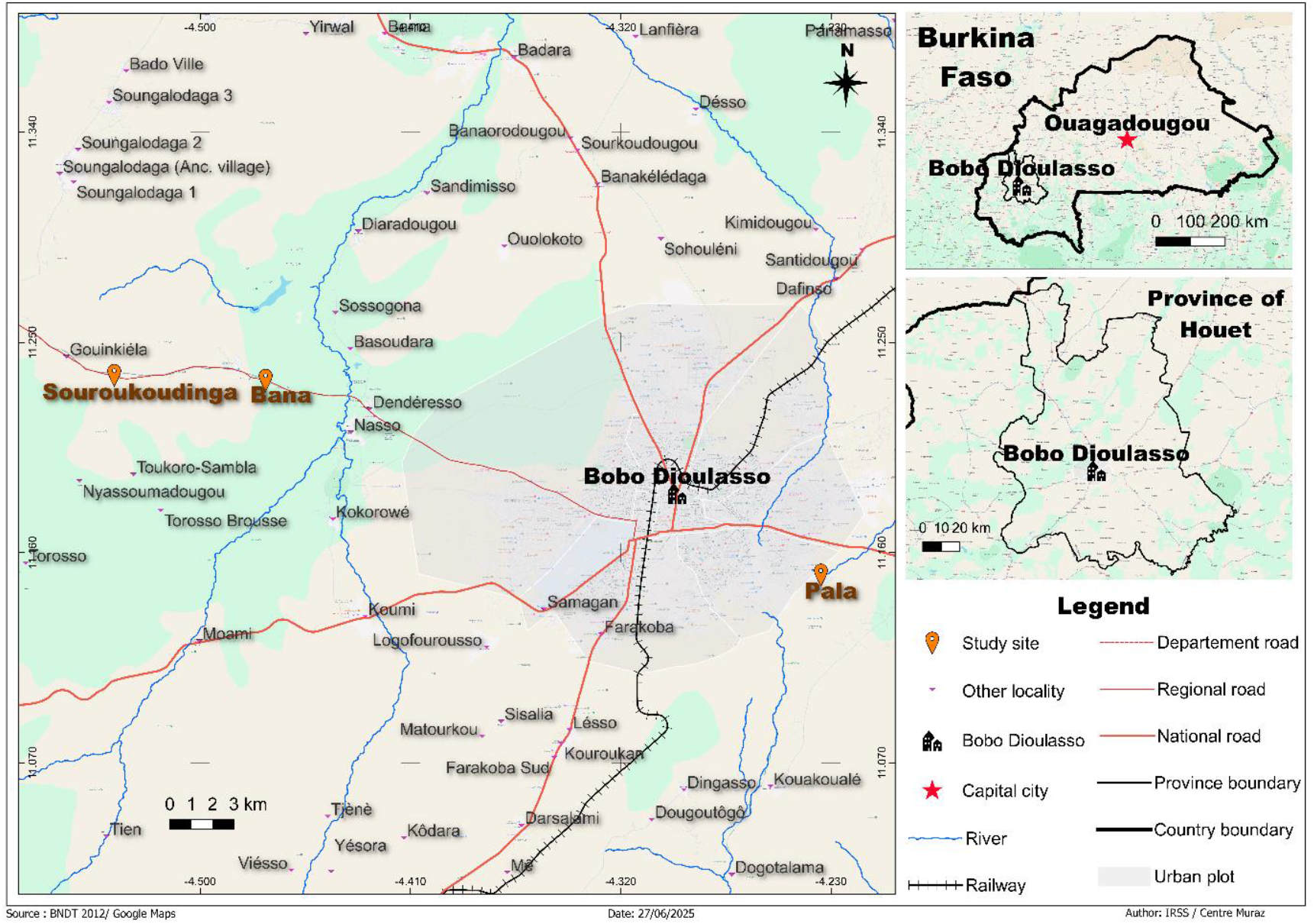
Location of our sampling locations in the Houet Province of western Burkina Faso. The coordinates of the sampling locations are: Bana [11.233, -4.472], Souroukoudingan [11.235, -4.537], Pala [11.151, -4.234]. In the centre of the map is Bobo-Dioulasso, the second largest city in Burkina Faso, where our research base locates. There are two sampling locations from Bana: Bana Village and Bana Market, which are about 23km from Bobo-Dioulasso. Souroukoudingan is about 7km further away in the same direction. Pala is on the opposite direction approximately 6km from Bobo-Dioulasso.

**Figure 2.**
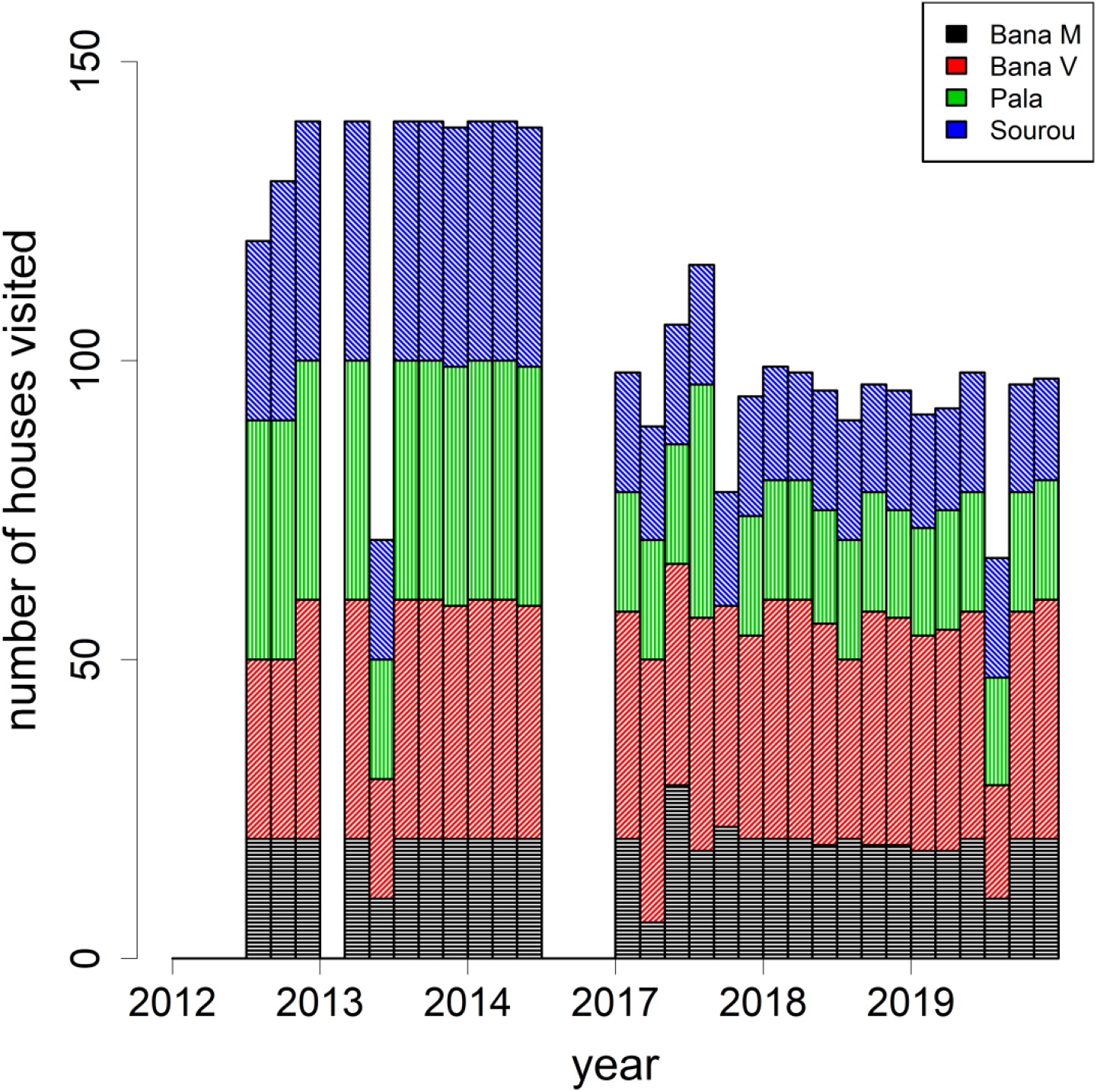
Sampling effort per village over time. Each bar represents the number of houses visited for PSC collection per two months. Sampling was paused from July 2015 to Dec 2016.

**Figure 3.**
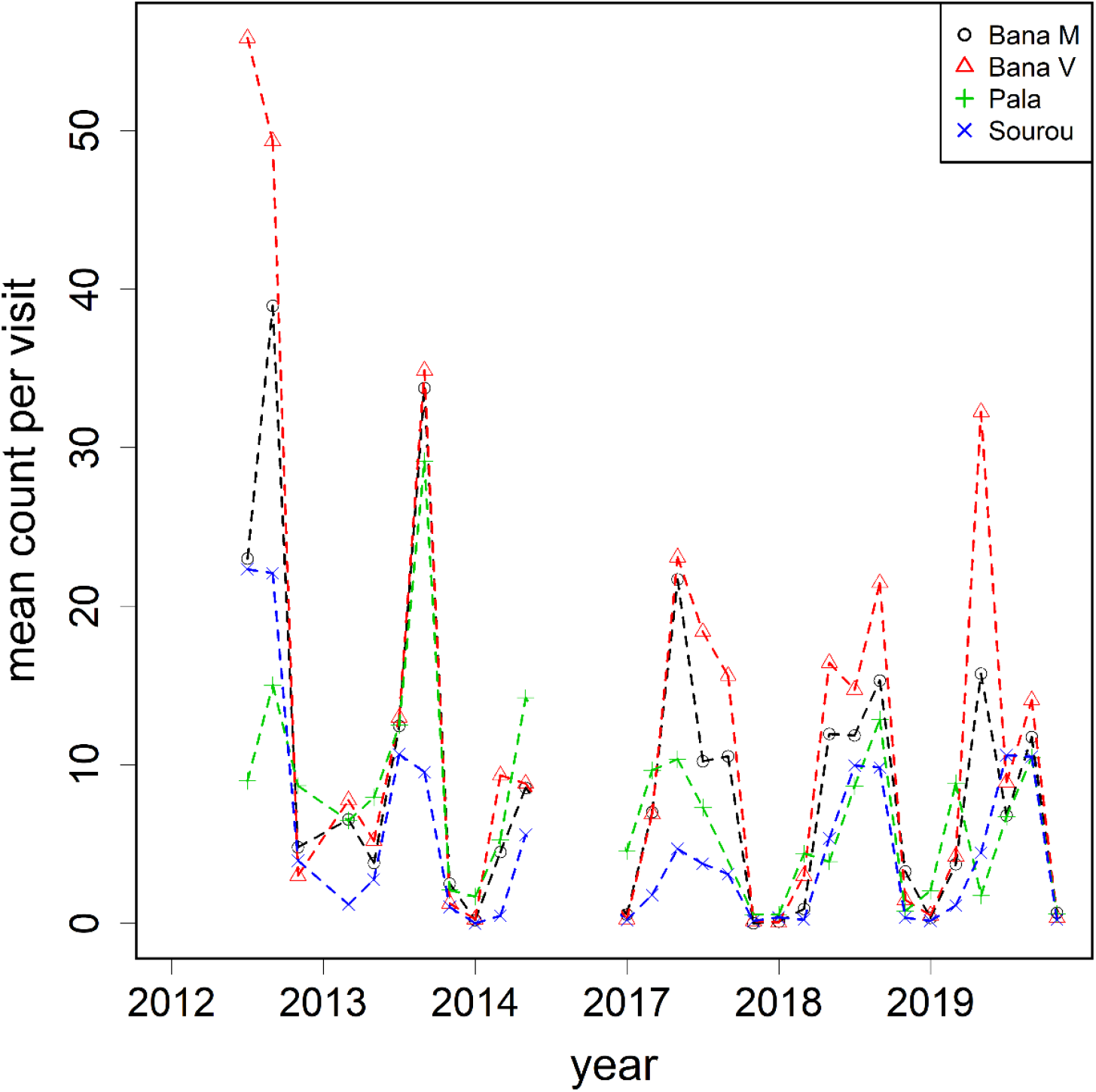
Mean female *An. gambiae s.l*. collected per visit by village over time. Note that sampling was paused after June 2014 then resumed at the beginning of 2017.

## Methods

Each house visit generated a line of record, including the number of female *An. gambiae s.l*., the date of collection, and a unique house ID. Other potential covariates concerning the visit were also documented, such as the number of residents, and whether bed net was in use. Female *An. gambiae s.l*. per visit was the main response to proxy indoor mosquito density. To model seasonality we used a six-level factor variable, with two consecutive months being pooled as one (e.g. January and February as one level, and so on) for a more balanced design. A Poisson generalised linear mixed model (GLMM) was fitted. Seasonality, household size, and the use of bed nets (yes/no) were treated as fixed effects. Observational random effects were included as a measure of overdispersion [18]. Villages, year of sampling, and unique house IDs were considered as random effects. The interaction between village and seasonality was also included for additional variability. For species composition, a similar binomial GLMM was fitted to the proportion between the female counts of the two most dominant species in the region. Analyses were conducted in R-4.3.1 [19] with packages lme4 and emmeans [20].

## Results

We endeavoured to follow closely the intended sampling plan with 3212 visits made (Figure 2). After removing entries with missing information, 3133 visits of 383 distinct houses were retained over the sampling period. Mosquito density as measured by the number of female *An. gambiae s.l*. is summarised as follows: A total of 28,919 female *An. gambiae s.l*. were collected, with an average of 9.2 per visit. The high mean count came with huge variation across visits ranging from 0 to 333 per visit. The mean household size was just over 3 persons – this value was subsequently used to classify household size into categories (none, small with 1-3 persons, large with >3 persons). Bed nets were installed in about 90% of the visits.

A Poisson GLMM was fitted to the counts (Table 1A). Seasonality was well defined within a calendar year with a period of low and high counts. Lower counts were observed between November and February, then increased when transitioning into the next season towards the peak at the months of September and October. There was about a 30-fold difference between the lowest and highest mean counts. As we grouped seasonality into six two-monthly factor levels, we performed contrasts to test whether their mean counts were different. All pairwise comparisons were significant, apart from the three levels of months between May and October where they shared the same mean. The presence of bed nets reduced mosquito count by 14%, but the effect was not statistically significant (p=0.099). Count increased with household size: compared to an empty house a 3-fold (p=0.013) increase was observed if there were ≤3 occupants, then a further 53% increase (p<0.001) for a bigger household with >3 occupants. For the random components (Table 1B), observation-level overdispersion had the highest contribution, accounting for 65% of the total variance. The second largest term was the between-house variation (14%). The between-village variance was found to be the lowest at 4.6%. The village-seasonality interaction allowing for additional variation on top of the two random components was estimated to be of moderate strength (8%). All five variance components were statistically larger than zero.

**Table 1A.** Fixed effect estimates from the Poisson GLMM on the counts of female *An. gambiae s.l*.. The significance of the fixed effects and p-values were based on Likelihood Ratio Tests (LRTs), while Wald p-values were computed for pairwise comparisons. Effect estimates are on log-scale. Asterisks are given to indicate statistical significance: (^***^) p<0.001, (^**^) p<0.01, (^*^) p<0.05.

| Fixed effect | df | Comparison(s) | Estimate (s.e.) | p-value |
| --- | --- | --- | --- | --- |
| Seasonality | 5 |  |  | 0.000*** |
|  |  | (See Table 1C) |  |  |
| Bed net | 1 |  |  |  |
|  |  | No vs Yes | -0.15072 (0.091) | 0.099 |
| Household size | 2 |  |  | 0.000*** |
|  |  | None vs Small | 1.110 (0.446) | 0.013* |
|  |  | Small vs Large | 0.427 (0.066) | <0.001*** |
|  |  | None vs Large | 1.537 (0.449) | 0.000*** |

**Table 1B.** Random effect estimates and percentage contributions from the same Poisson GLMM, ranking from the largest. LRTs were performed to obtain the p-values.

| Random effect | Number of levels | Variance estimate | Percentage | p-value |
| --- | --- | --- | --- | --- |
| Overdispersion | 3133 | 1.4413 | 64.88 | 0.000*** |
| Between-house | 383 | 0.3127 | 14.07 | 0.000*** |
| Between-year | 6 | 0.1881 | 8.47 | 0.000*** |
| Between-village x<br>Seasonality | 24 | 0.1768 | 7.96 | 0.000*** |
| Between-village | 4 | 0.1025 | 4.61 | 0.048* |

**Table 1C.** Pairwise contrast for the seasonality effect in the Poisson GLMM (Table 1A). Seasonality was categorised into 6 factor levels. For example, “0102” refers to the counts of Jan and Feb combined. Contrasts (lower-triangular) and the unadjusted p-values (upper-triangular, shaded in grey) were computed via the R package “emmeans”.

|  |  |  |  |  |  |  |
| --- | --- | --- | --- | --- | --- | --- |
|  | 0102 | 0304 | 0506 | 0708 | 0910 | 1112 |
| 0102 |  | 0.000*** | 0.000*** | 0.000*** | 0.000*** | 0.000*** |
| 0304 | 2.151 |  | <0.001*** | 0.003** | 0.000*** | 0.000*** |
| 0506 | 3.212 | 1.061 |  | 0.638 | 0.285 | 0.000*** |
| 0708 | 3.066 | 0.915 | -0.146 |  | 0.130 | 0.000*** |
| 0910 | 3.544 | 1.394 | 0.332 | 0.479 |  | 0.000*** |
| 1112 | 0.798 | -1.353 | -2.414 | -2.268 | -2.746 |  |

A subset of female *An. gambiae s.l*. from the second phase of collection (2017-2019) was genotyped for species identification. After genotyping, 3626 were identified as *An. coluzzii* and 1102 as *An. gambiae*, yielding an overall ratio of approximately 3:1 from 985 house visits (Figures 5-6). Apart from the two dominant species a small proportion of *An. arabiensis* (∼5%) was also found but excluded from subsequent statistical analyses. The species proportions in our four villages were very different (Figure 4). In Bana Market and Bana Village *An. coluzzii* was the dominant species making up over 90% of the samples, and similarly 60% in Souroukoudingan, while in Pala it was the minority (24%). Although species composition varied throughout a year, the four villages did not seem to follow the same dynamics nor a pre-defined pattern. A binomial GLMM was then fitted to model the relative proportion of the two species. For the fixed effects, neither the use of bed nets (p=0.77) nor household size (p=0.89) had an influence on the proportion (Table 2A). For the random components (Table 2B), the between-village variation was estimated to be the largest, explaining 77% of the total random variance. This was followed by overdispersion (11.4%) and the village-seasonality interaction (3.9%). The between-house variation was not significant (1.5%, p=0.26). In other words, setting overdispersion aside, the relative importance of the random components were the exact opposite of those for mosquito density.

**Table 2A.**
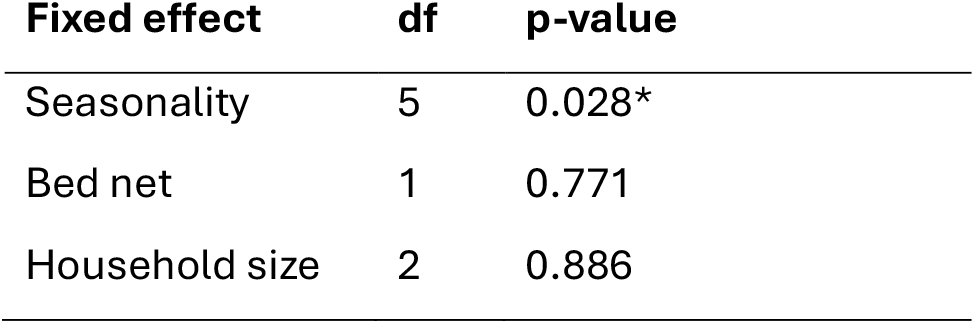
Fixed effect estimates from the binomial GLMM on the proportion between female *An. coluzzii* and *An. gambiae*, with p-values obtained from LRTs. Effects are reported on logit scale.

| Fixed effect | df | p-value |
| --- | --- | --- |
| Seasonality | 5 | 0.028* |
| Bed net | 1 | 0.771 |
| Household size | 2 | 0.886 |

**Table 2B.** Random effect estimates and percentage contributions from the binomial GLMM on species proportion, ranking from the largest. LRTs were performed to obtain the p-values.

| Random effect | Number of levels | Variance estimate | Percentage | p-value |
| --- | --- | --- | --- | --- |
| Between-village | 4 | 3.476 | 76.47 | 0.000*** |
| Overdispersion | 985 | 0.520 | 11.43 | 0.000*** |
| Between-village x Seasonality | 24 | 0.304 | 6.68 | 0.000*** |
| Between-year | 3 | 0.179 | 3.92 | 0.000*** |
| Between-house | 200 | 0.0678 | 1.49 | 0.263 |

**Figure 4.**
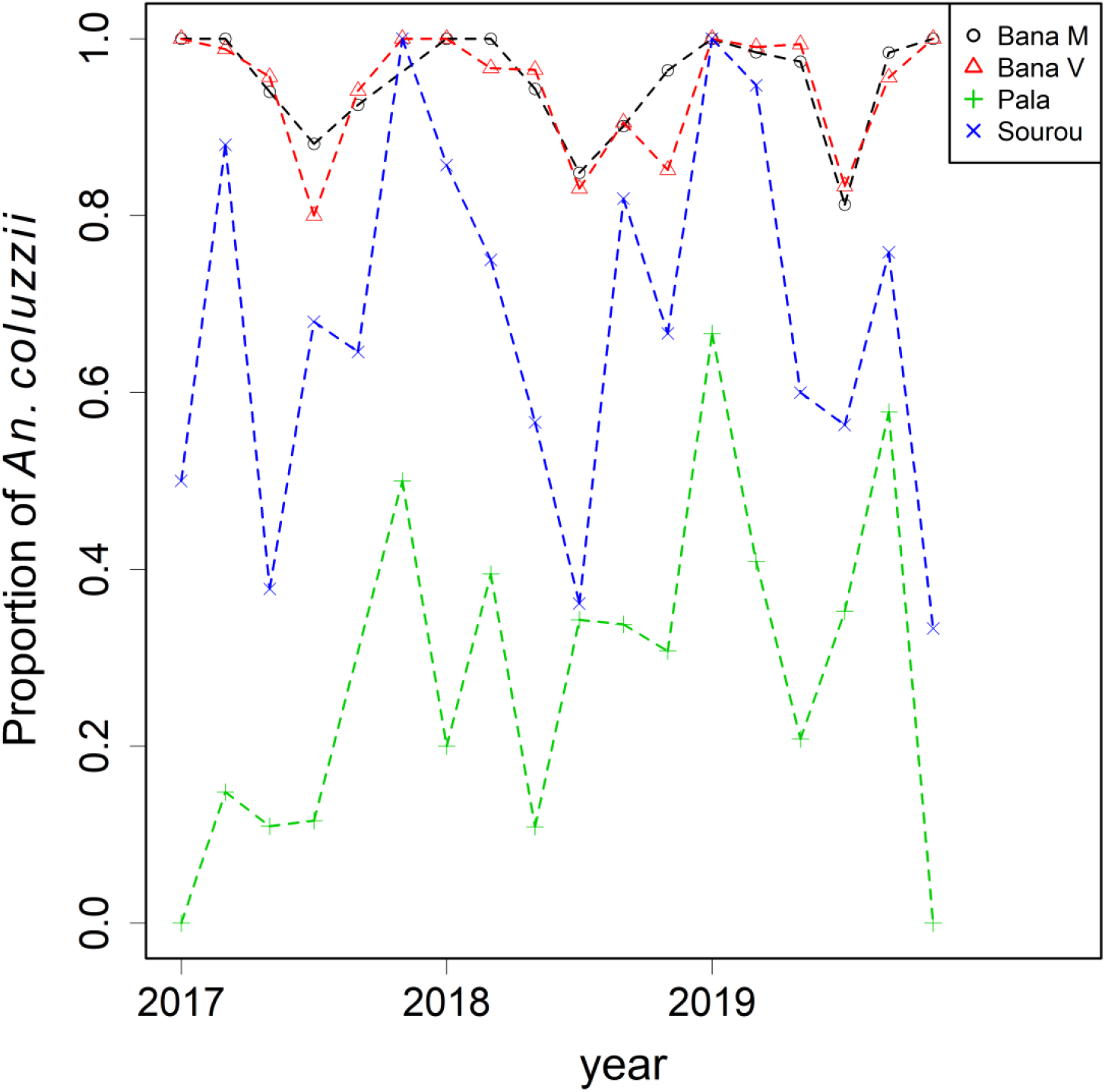
The relative proportion of *An. coluzzii* and *An. gambiae* by village over time. Species identification was performed to a subset of mosquitoes collected during 2017-2019.

**Figure 5.**
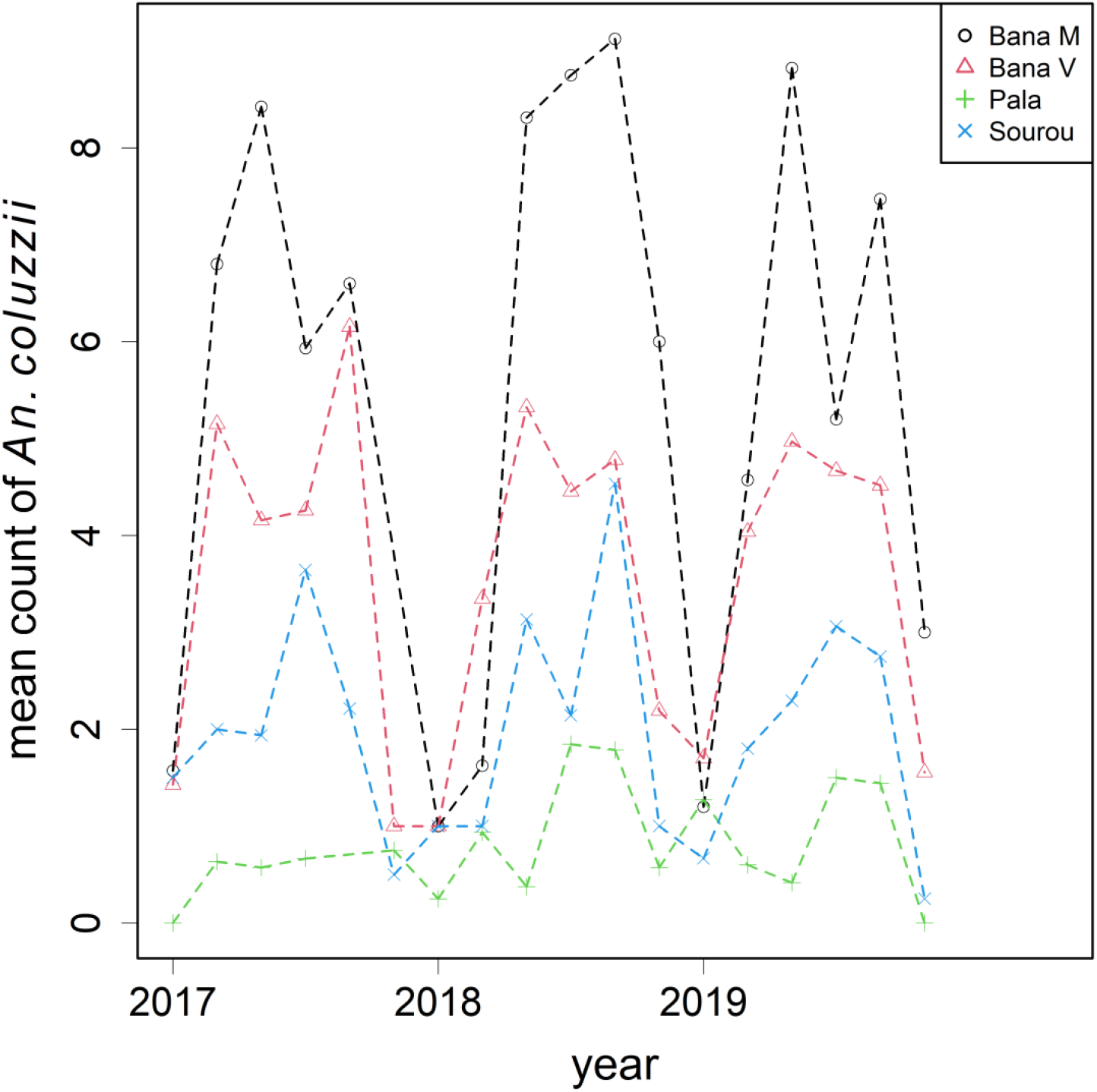
Mean number of female *An. coluzzii* per house visit by village over time.

**Figure 6.**
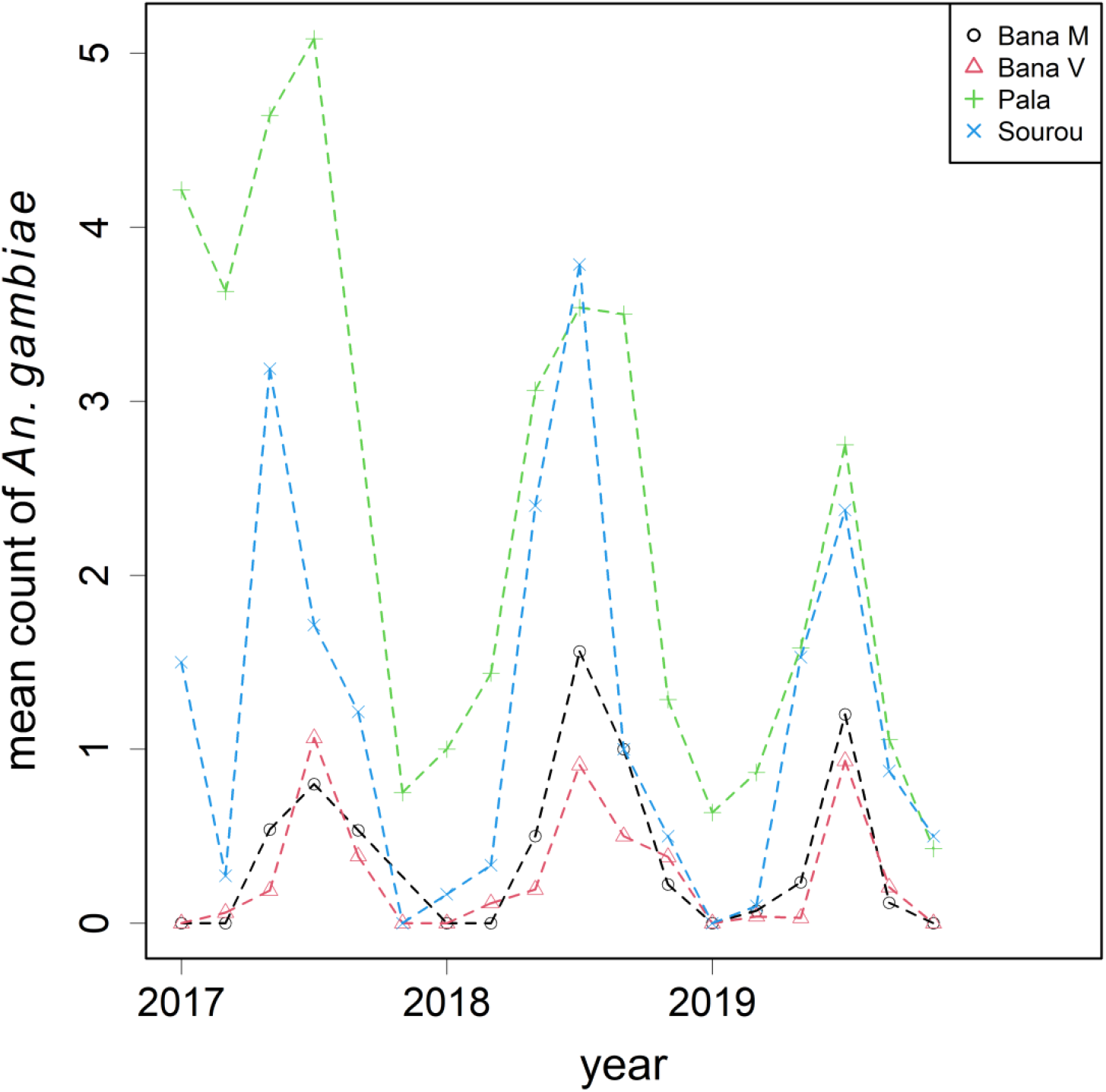
Mean number of female *An. gambiae* per house visit by village over time.

## Discussion

Several baseline collections have been conducted in Burkina Faso and beyond to evaluate vector density over space and time [21-23]. Our results agree with the existing knowledge that mosquito density (as measured by female counts in this study) exhibits strong seasonal trends. The study region is characterised by a tropical climate marked by two main seasons, with average annual temperatures between 25-30°C and rainfall of 800-1100mm [24]. The rainy season is usually defined as the months from June to September, with May and October being in transition [15], the same period at which we consistently observed higher counts. The peak at the late rainy season near October was also detected by Dao et al. [12] in a Sahelian village in the neighbouring country of Mali. After reaching the peak a period of low counts immediately followed from November into February. The clean switch between the two seasons is a signature of the Sahelian region as the surface water disappears, drastically reducing the availability of larval sites [25, 26]. In parallel to our census size estimates, genetic estimates of effective population size also showed similar patterns of seasonal variation [27].

While many existing collections aimed to correlate seasonality and other deterministic drivers with mosquito dynamics through monthly summaries, curating individual PSC records gives opportunity to analyse covariates on finer scales such as on observational and household level. Bed net use, a factor considered as a fixed effect in this study, only reduced mosquito density by 14% (but not significant) despite a high coverage of over 90%. A separate study discovered the increase in frequency of insecticide resistance alleles during the same period which potentially explains its relatively low efficacy [3]. We found a strong relationship between density and household size, a factor which was previously thought to be less important [21]. Another dimension of this work is to estimate the random effects, which induce uncontrolled heterogeneity that have not been explained by our main effects. We intentionally fitted a 6-level seasonality variable with 24 additional random village-seasonality interaction terms to our GLMMs to saturate the seasonality effect, allowing us to focus on these random components. The sizes and relative importance of the random components are of interest rather than how and why they differ. Several random components on various levels were considered: Overdispersion, assigned to each visit, was also the largest component on mosquito density. It includes the intrinsic variation of indoor mosquito density, sampling method, and potentially other abiotic or idiosyncratic factors. By revisiting some households, the between-house variance was estimated to be the second largest component, hinting some houses were more susceptible to mosquitoes even though they were exposed to similar conditions as their neighbours. The difference could be due to building material, usage, and proximity to larval sites. Further modelling, including the use of GIS data, is warranted to further dissect this variance component [21]. The between-village and between-year variances were minor with respect to density, each contributing to only a few percent of the total random variation.

There is no surprise that the two primary vectors *An. coluzzii* and *An. gambiae* were abundant in the region. Our binomial GLMM concluded that the proportion of the two species was unaffected by bed net usage or household size. The four sampled villages tended to have their own underlying proportions and seasonal variations: Souroukoudingan and the two sites from Bana were predominantly *An. coluzzii*, while Pala was the opposite. There was no observable switch in the proportions across seasons. The large between-village and village-seasonality interaction illustrated how diverse species composition can be across landscape. It is known that the two species coexist but with rather contrasting preferences. *An. coluzzii* prefers sites with permanent freshwater while *An. gambiae* can tolerate intermittent access to water [25]. Bana and Souroukoudingan have access to water year-round due to agricultural activities that explains the strong presence of *An. coluzzii*. Their respective persistence mechanisms may also affect the relative abundance in the dry season and transition months. It is generally thought that *An. coluzzii* aestivates locally during the dry season thus an immediate population expansion is expected when the aestivators reemerge, whereas *An. gambiae* takes days to respond to the arrival of the rainy season as immigrants from further afield recolonise the focal population [28]. In short, the large between-village and lack of between-house variance were the key signatures of species composition. In fact, the relative importance of the random components in species composition is the exact opposite of that of density (Table 3), which itself is a very interesting finding. The design of population monitoring programmes and control strategies will therefore be influenced by the choice of response variable (see below). Future work should include the monitoring of *An. arabiensis*, which was also present in our study, especially in urban settings where it appears to be colonising [29].

**Table 3.**
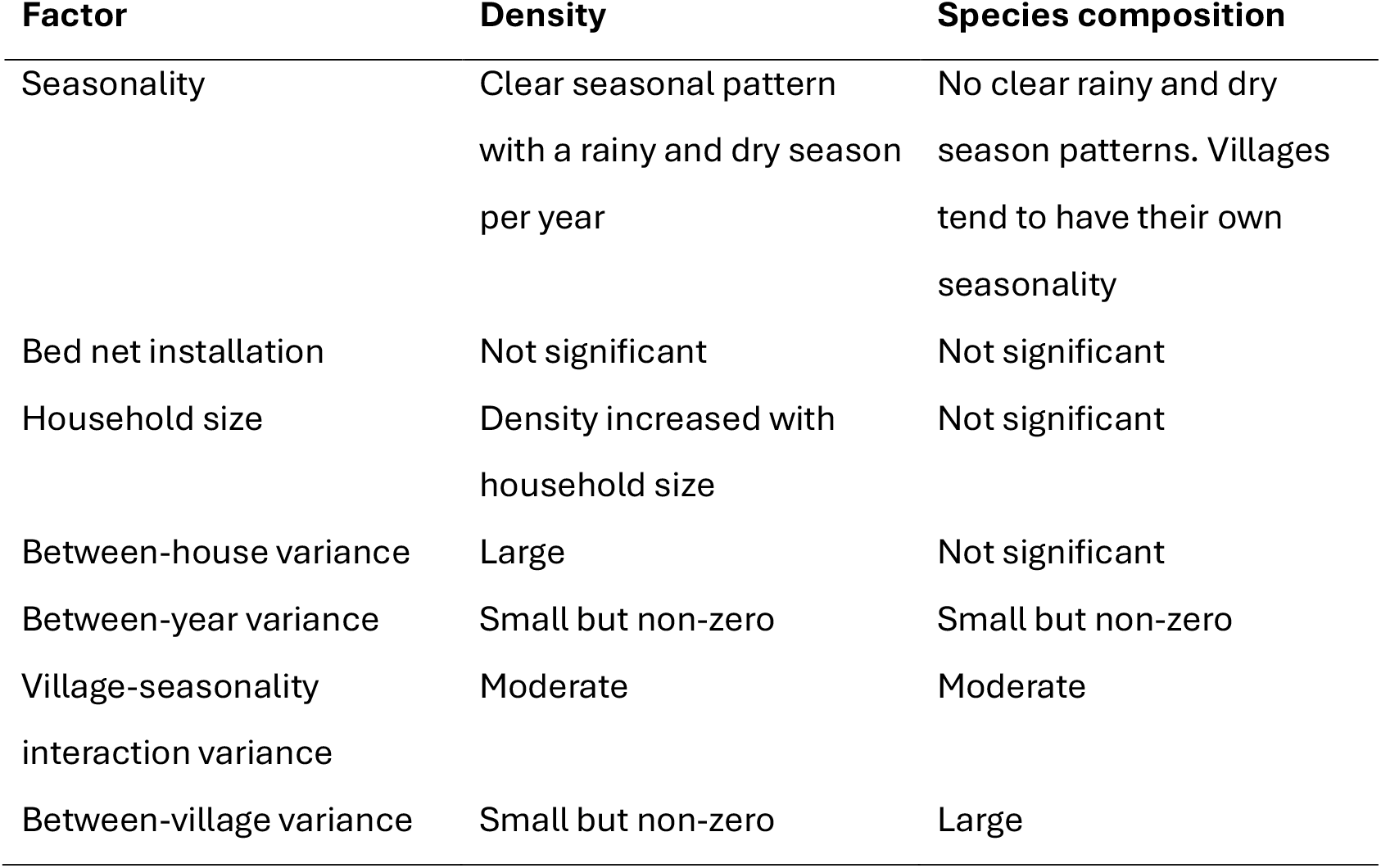
Descriptive summary of the fixed and random factors affecting mosquito density and species composition, based on the numerical results from the two GLMMs reported in Tables 1-2. For the random effects, the relative importance is reported.

Both mosquito density and malaria transmission are subject to strong spatial-temporal variation. Besides our findings on density, a previous survey in the same region estimated a 25-fold increase in EIR from the dry to rainy season, mainly driven by the sharp rise in human landing rate [22]. Both malaria incidence and mosquito density found their peaks towards the late rainy season in Burkina Faso [30], meaning seasonal strategies may be required [31], further straining the healthcare system. Seasonality may also affect the overall efficacy of GM vector control and the optimal schedule of releases [32, 33]. While the heterogeneity in transmission across seasons or climatic zones have been anticipated, the unexplained variance components impose additional challenges on the road to the elimination of malaria. Persistent transmission can be maintained even under continued and effective malaria control [34]. It only requires a minority of human individuals with dry season exposure to constitute an infectious reservoir [35]. The higher the random variances in mosquito density the higher chance these small pockets of space are created for permanent transmission. Classical epidemiological models showed that with heterogeneous biting, a factor of (1 + α) is applied to *R*_0_ where α is the squared coefficient of variation (CV) in biting rate. In other words, *R*_0_ increases linearly with the variance in biting rate if all else is equal. While biting rates are best estimated from human landing catches, our PSC collection and between-house variance estimate in density could be carefully extrapolated to inform the degree of heterogeneity linking to indoor exposure and biting. With the many layers of random variations, the compounding effect on *R*_0_ could potentially be significant even if the mean estimate suggests otherwise. Biting can also occur outdoor, thus measuring the outdoor mosquito density and species composition with other trapping methods would complement this study [36]. In terms of reporting, increased spatial and temporal heterogeneity in malaria cases in low transmission settings reduces the usefulness of national or regional level trends in incidence or prevalence [37].

Any new intervention needs to undergo phases of rigorous testing and trial. A cRCT is the gold-standard study design for evaluating public health programmes, where interventions are addressed to some clusters then compared to those without [38, 39]. Our entomological dataset and parameter estimates can contribute towards baseline data collection for a potential upcoming field trial, and more importantly, inform some of the key considerations concerning trial design. In many mosquito intervention programmes treatments must be addressed to a cluster of households (e.g. a village) as a whole, hence it is the basic unit for randomisation. If the proposed intervention targets the entire species complex to suppress mosquito density by a certain percentage, then its effect can be evaluated from the difference in counts between the control and treatment arms via a Poisson GLMM like the one we have fitted but with an additional variable indicating the treatment arm. We suggest sampling should prioritise months with high abundance to maximise the chance of detecting the intervention effect. There are two reasons for this: The large baseline counts in the rainy seasons provide headroom for population suppression, while in the drier months the monthly aggregated counts are low with numerous zero readings from individual houses. The second reason is that the intervention effect is likely to be estimated with greater precision (i.e., a lower coefficient of variation). Observation-level covariates such as household size were shown to be correlated with the baseline counts. For the same reason, the recruitment of households should target those with habitants. These covariates are essential when the cRCT is analysed from observation-level data [40], and remain useful under the use of cluster-level summaries, where they can inform the pairing of clusters in restricted randomisation, or even a matched-pair cRCT design to further improve balance and statistical power [41].

The random effect estimates will also influence the design of the cRCT. The current sampling of 20 houses per month per village aimed to illustrate seasonality by averaging out the overly dispersed count data. One can find the optimal sampling strategy by altering the number of clusters and houses per month per cluster while keeping the total sampling effort constant. Given the relatively high importance of the between-house variation, sampling the same set of houses repeatedly may risk inducing systematic bias. Small but non-zero between-year variation in density was detected across the 6-year period. Yearly variation is less of a problem for the classical parallel cRCT design as comparisons are made mostly within the same time period (vertical) between the two arms. A stepped-wedged design, however, contains both vertical and within-cluster (horizontal) comparisons, may be prone to this effect. In the stepped-wedged design, clusters are randomised into several streams. All clusters are in control condition initially and will have received the intervention at the end, with intervention being gradually rolled out at different time points of the trial. As all clusters now include pre- and post-intervention phases, a good year (with respect to mosquitoes’ survival) alone may result in a sudden increase in mosquito density even in clusters receiving the intervention. On a positive note, the between-year variance was estimated to be small with limited impact, contributing to only a few percent of the total variation in both mosquito counts and species proportions. While our four villages were chosen based on logistical reasons, they were considered as a random effect in the GLMMs as this would be the case in a cRCT. A non-zero between-village variance was estimated despite them being in the close vicinity (∼30km radius) of Bobo-Dioulasso. A full-scale cRCT would likely be conducted on a regional or even national basis with many more clusters, thus this variance is expected to increase with geographical coverage. Note that these parameter estimates (Tables 1A, 1B) are reported in the log scale, which means they should be compared to the intervention strength of log(1-G) if the intervention suppresses the density by G. In trial designs, the between-village variance informs the intra-cluster correlation coefficient (ICC), which measures relative similarity of the observations from the same cluster compared to those from the others. Since individual variances (i.e. overdispersion and between-house variation) were estimated to be much larger than between-village variance, the ICC is expected to be reasonably small, which helps reduce the number of clusters required.

Some novel vector control strategies can be tailored to target one or a subset of species while keeping the rest intact. The results from such a cRCT can be analysed with a similar Poisson GLMM on the counts of the targeted species, potentially with the non-targeted species as covariates. Another way is to analyse the relative proportion of the two as a pair of joint response with a binomial GLMM, whose intervention effect is log(1-G) on the logit scale. Between *An. coluzzii* and *An. gambiae*, the ICC in proportion is inferred to be much higher than in their combined density. In other words, most variation comes from clusters rather than the individual houses, thus the effective sample size approaches the number of clusters. Under high ICC the stepped-wedged design may be preferred as the efficiency is improved by the inclusion of horizontal comparisons [42]. The lack of between-house variation in species composition has the potential to simplify the recruitment process, as one or a few households could represent the entire cluster. To conclude, our data and analyses provide crucial parameter estimates to be used in trial design, power analyses and sample size determination. With the many combinations of variables and scenarios, quantitative studies are necessary to fully explore their complex interplay, often aided by computer simulations.

## Author contributions

P.S.E., A.A.M., F.A.Y. led the field collection. P.S.E., D.K. conducted the genotyping. P.S.E., A.A.M., F.N., T-Y.J.H curated and processed the data. T-Y.J.H. conducted the statistical analyses. T-Y.J.H wrote the manuscript with inputs from the other authors.

## Conflicts of interest

Authors declare no conflicts of interest.

## Acknowledgements

We thank John Connolly, Samantha O’Loughlin, Silke Fuchs, and Camilla Beech for their feedback on an earlier draft. All authors receive funding through Target Malaria, which is supported by a grant from the Bill & Melinda Gates Foundation and from the Open Philanthropy Project Fund, an advised fund of the Silicon Valley Community Foundation. This work is supported by Wellcome Trust (224487).

## Data availability

The dataset will be made available upon acceptance.

## Notes

### Competing Interest Statement

The authors have declared no competing interest.

### Summary of Updates

Author list updated, plus major revision.

